# Clonal heterogeneity influences the fate of new adaptive mutations

**DOI:** 10.1101/039859

**Authors:** Ignacio Vázquez-García, Francisco Salinas, Jing Li, Andrej Fischer, Benjamin Barré, Johan Hallin, Anders Bergström, Elisa Alonso-Perez, Jonas Warringer, Ville Mustonen, Gianni Liti

## Abstract

**In Brief:** Vázquez-García et al. examine the role of clonal heterogeneity in the acquisition of antimicrobial resistance. They report that pre-existing and *de novo* genetic variation jointly contribute to clonal evolution. By building a library of adaptive mutations in multiple genetic backgrounds, they resolve the fitness effects of mutations in a clonal lineage.

**Highlights:**

- Clonal heterogeneity influences the acquisition of antimicrobial resistance
- Joint role of pre-existing and *de novo* genetic variation in clonal evolution
- Clonal dynamics are shaped by background-dependent fitness effects of mutations
- Loss of clonal heterogeneity is balanced by genomic instability and diversification

**Summary:** The joint contribution of pre-existing and *de novo* genetic variation to clonal adaptation is poorly understood, but essential to design successful antimicrobial or cancer therapies. To address this, we evolve genetically diverse populations of budding yeast, *S. cerevisiae*, consisting of diploid cells with unique haplotype combinations. We study the asexual evolution of these populations under selective inhibition with chemotherapeutic drugs by time-resolved whole-genome sequencing and phenotyping. All populations undergo clonal expansions driven by *de novo* mutations, but remain genetically and phenotypically diverse. The clones exhibit widespread genomic instability, rendering recessive *de novo* mutations homozygous and refining pre-existing variation. Finally, we decompose the fitness contributions of pre-existing and *de novo* mutations by creating a large recombinant library of adaptive mutations in an ensemble of genetic backgrounds. Both pre-existing and *de novo* mutations substantially contribute to fitness, and the relative fitness of pre-existing variants sets a selective threshold for new adaptive mutations.

## Introduction

The adaptive response of a cell population can thwart therapeutic control of a wide spectrum of diseases, from bacterial and viral infections to cancer. A prototypical scenario arises when individuals in a population acquire heritable genetic or non-genetic changes to adapt and thrive in a new environment (Balaban et al., 2004; Marusyk et al., 2014; Toprak et al., 2012). Since the seminal findings by Luria and Delbrück that phage-resistant bacteria can acquire adaptive mutations prior to selection (Luria and Delbrück, 1943), measuring the fitness effects and dynamics of mutations has been key to map the principles of evolutionary adaptation (Barrick and Lenski, 2013). The focus has typically been on characterizing few mutations at a time under the implicit assumption that beneficial mutations are rare, treating pre-existing and acquired mutations separately. However, many mutations are often simultaneously present in a population, which result in fitness differences between individuals that selection can act upon (Lang et al., 2013; Levy et al., 2015; Parts et al., 2011; Venkataram et al., 2016).

Given that mutations in asexual populations are physically linked in the genome, the fates of preexisting and *de novo* mutations are mutually dependent and selection can only act on these sets of variants in their entirety. Genome evolution experiments on isogenic populations have revealed both adaptive sweeps and pervasive clonal competition in large populations where the mutation supply is high. This phenomenon, known as clonal interference, takes place as mutations in different individuals cannot recombine via sexual reproduction and is now relatively well understood both experimentally and theoretically (Gerrish and Lenski, 1998; Lang et al., 2013; Neher, 2013). Experiments on populations with extensive genetic variation have demonstrated that beneficial mutations expand in a repeatable way (Parts et al., 2011). Theory predicts that the rate of adaptation is proportional to the fitness variance present in a population, generating a traveling fitness wave (Desai and Fisher, 2007; Rouzine and Coffin, 2005). However, the role of *de novo* mutations has been negligible in these experiments, either because of their short duration or related to the selective constraints used. A study which was able to anticipate new mutations found that one or few genetic variants were sufficient to affect the fate of subsequent beneficial mutations, hinting that the joint dynamics of new mutations have to be considered in the light of pre-existing variation (Lang et al., 2011). The ensuing interaction between existing and subsequent mutations has been theoretically considered under different population genetic scenarios (Good et al., 2012; Hermisson and Pennings, 2005; Orr and Betancourt, 2001; Peter et al., 2012; Schiffels et al., 2011). A key theoretical prediction is that a new beneficial mutation will only establish if it has a selective advantage greater than a characteristic value that depends on the underlying fitness distribution (Good et al., 2012; Schiffels et al., 2011). However, this important hypothesis remains to be tested: namely, whether genetic diversity can change the evolutionary fate of new adaptive mutations by limiting the number of backgrounds where they can still outcompete the fittest extant individuals. Understanding the impact of genetic heterogeneity on adaptive dynamics is particularly urgent as recent findings indicate that it can play a major role in the development of resistant bacterial infections (Lieberman et al., 2014) and in cancer recurrence (Gerlinger et al., 2012; Landau et al., 2013).

We have delineated two lines of enquiry into our hypothesis: (i) To what extent can the adaptive response be attributed to genetic variation already present in a population and how much to acquired? (ii) How do the aggregate effects of pre-existing variation influence the fate of new mutations? To address these questions, we investigated the interaction between pre-existing (or background) genetic variation and new mutations in a population of diploid cells with unique combinations of alleles. The cells originate from two diverged *S. cerevisiae* strains (Figure 1). We carried out 12 rounds of random mating and sporulation (meiosis) between DBVPG6044, a West African palm wine strain (WA), and YPS128, a North American oak tree bark strain (NA) (Parts et al., 2011). The cross population (WAxNA) consisted of 10^7^-10^8^ unique haplotypes, with a pre-existing single-nucleotide variant segregating every 230 bp on average. We further identified 91 *de novo* single-nucleotide variants (SNVs) and small insertions and deletions (indels) acquired during the crossing phase from genome sequences of 173 founder individuals. This is consistent with a mutation rate of approximately 2.89×10^−10^ mutations per base per generation, close to empirical estimates in other yeast strains (Zhu et al., 2014). We also observed aneuploidy in chromosome IX, indicating the presence of variation other than point mutations. This design results in the frequency spectrum of background mutations to be normally distributed, so that pre-existing variants are already established and do not need to overcome genetic drift. We refer to the parental genotype of each individual in the cross as its genetic background, which on average differ by ~31,000 SNVs between individuals. Since naturally occurring deleterious mutations have been selected against over long evolutionary timescales, the recombinant parental genotypes are enriched for functional diversity which is not readily accessible using other techniques, such as random or site-directed mutagenesis. The cross-based approach also reduces genetic linkage of nearby loci, which enables us to localize background alleles responding to selection.

**Figure 1:**
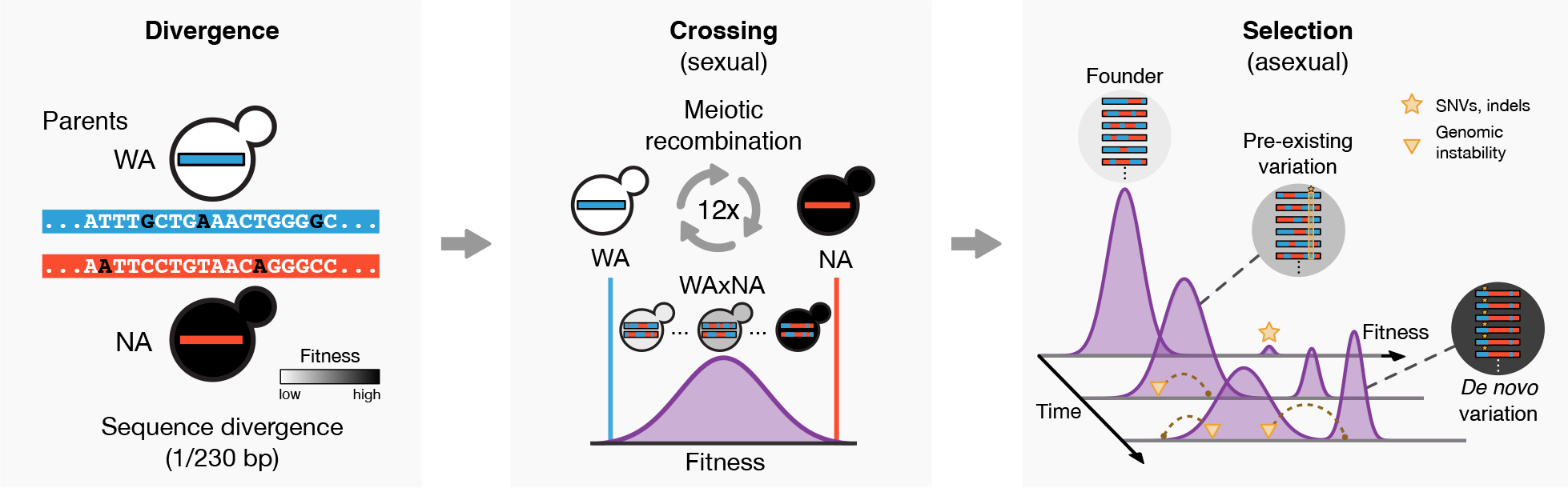
Study Overview. Schematic diagram of the divergence, crossing and selection phases of the experiment. Two diverged *S. cerevisiae* lineages (WA and NA) were crossed for twelve rounds, generating a large ancestral population of unique haplotypes. These diploid cells were asexually evolved for 32 days in stress and control environments and their adaptation was studied by whole-population and isolate sequencing and phenotyping. Populations evolved resistant macroscopic subclones driven by individual cells with beneficial genetic backgrounds (i.e., parental allele configurations) and by beneficial *de novo* mutations which provided a resistance phenotype.

Starting from WA, NA and WAxNA founders, we asexually evolved populations of ~10^7^ cells in serial batch culture under drug inhibition with hydroxyurea (HU) and rapamycin (RM) at concentrations impeding, but not ending, cell proliferation. These drugs were chosen for having known targets and in order to cover two of the most common modes of action of antimicrobial and chemotherapy drugs: inhibition of nucleic acid synthesis (hydroxyurea) and inhibition of protein synthesis and cell growth (rapamycin). We derived replicate lines of WA, NA (2 each in hydroxyurea and rapamycin) and WAxNA (6 in hydroxyurea, 8 in rapamycin and 4 in a control environment), propagating them for 32 days in 48-hour cycles (~54 generations; Experimental Procedures). We monitored evolutionary changes by whole-genome sequencing of populations after 2, 4, 8, 16 and 32 days, as well as clonal isolates at 0 and 32 days (Table S1). Finally, we measured the rate of growth at the initial and final time points for a subset of populations, and quantified the relative fitness contributions of background and *de novo* variation using a genetic cross.

## Results

Two regimes of selection became readily apparent in both sequence and phenotype. Initially, there were local changes in the frequency of parental alleles under selection (Figure 2). Over time, subclonal populations arose and expanded, depleting the pool of genetic diversity. Here and throughout this Article, we employ the term ‘subclone’ to refer to a group of cells that carry the same set of mutations. These successful ‘macroscopic’ could be detected by whole-population sequencing and phenotyping, persisting in time as manifested by broad jumps in the allele frequency visible across the genome and by multiple modes in the fitness distribution (Figure 2, Figure 3 and S2). But what drives these clonal expansions: is it the founder haplotypes themselves, *de novo* mutations relegating the parental variation to the role of passengers, or their combined action?

**Figure 2:**
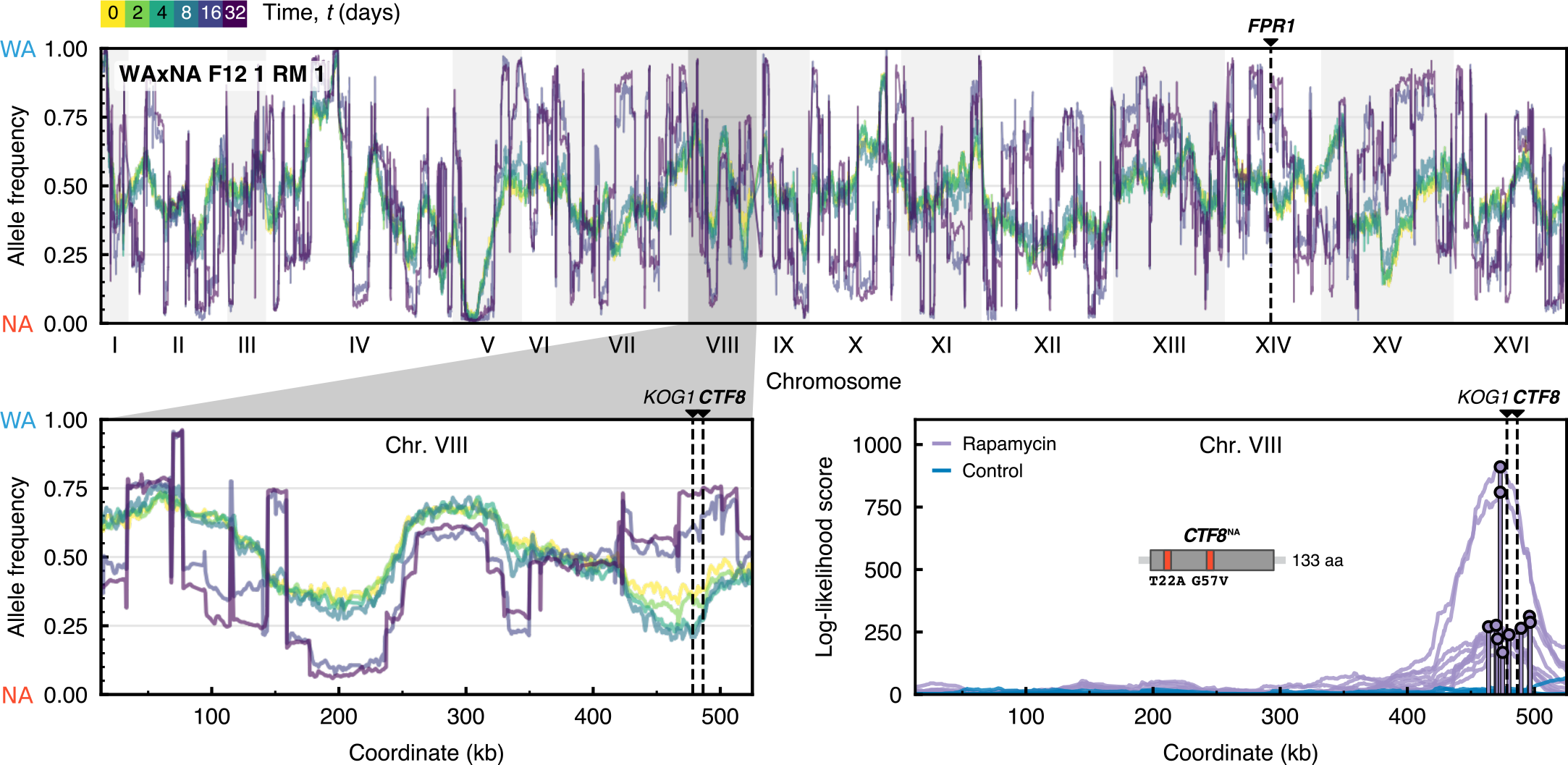
Genome-Wide Allele Frequency Changes. Genome-wide allele frequency of pre-existing parental variants after *t* = (0,2,4,8,16,32) days, measured by whole-population sequencing for a representative population in rapamycin. Pre-existing and *de novo* driver mutations are highlighted by dashed lines. Top panel: Chromosomes are shown on the *x*-axis; the frequency of the WA allele at locus *i*, 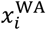, is shown on the *y*-axis. The reciprocal frequency of the NA allele is equivalent since 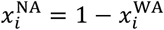. Bottom left panel: Zoomed inset of the shaded region shows allele frequency changes in chromosome VIII during selection in rapamycin. Early time points 2, 4 and 8 show localized allele frequency changes at 460-490 kb due to a beneficial NA allele sweeping with hitchhiking passengers. Late time points 16 and 32 show abrupt jumps between successive loci that reflect the parental haplotype of emerging subclone(s). These long-range correlations can alter the frequency of parental alleles independently of their fitness value. In case of a fully clonal population, allele frequencies at 0, 0.5 and 1.0 would correspond to the background genotypes NA/NA, WA/NA, and WA/WA of a diploid clone that reached fixation. Bottom right panel: We tested a model where each allele is proposed to be a driver under selection, with linked passenger alleles also changing in frequency by genetic hitchhiking. Top log-likelihood scores are shown for all populations in this region of interest (Supplemental Experimental Procedures). We validated the *CTF8*^NA^ allele to be strongly beneficial for rapamycin resistance (Figure S8). See also Figures S1 and S2.

**Figure 3:**
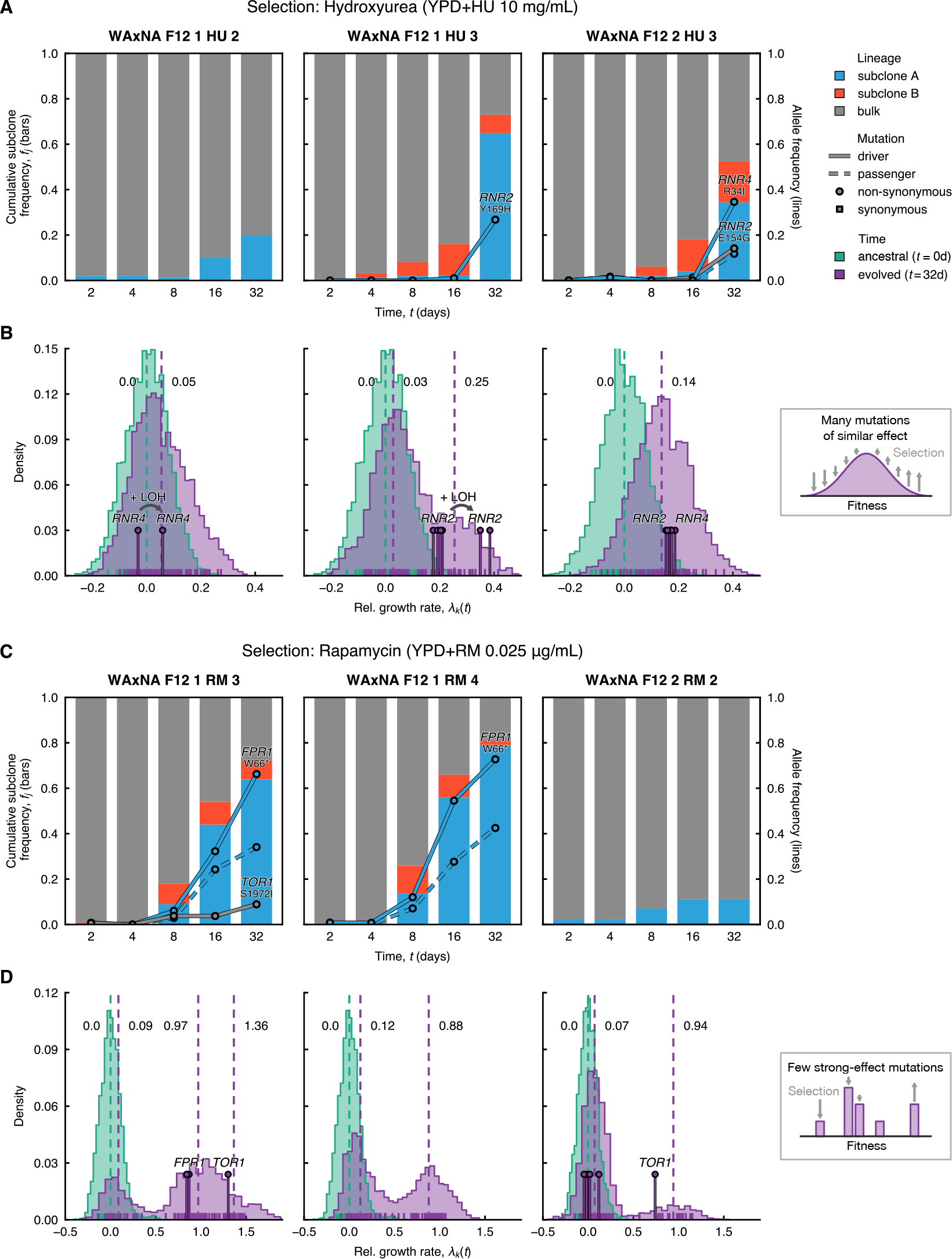
Reconstruction of Subclonal Dynamics. Competing subclones evolved in hydroxyurea and rapamycin experienced a variety of fates. (A and C) Time is on the *x*-axis, starting after crossing when the population has no macroscopic subclones. Cumulative haplotype frequency of subclones (bars) and allele frequency of *de novo* mutants (lines) are on the *y*-axis. Most commonly, selective sweeps were observed where a spontaneous mutation arose and increased in frequency. Driver mutations are solid lines and passenger mutations are dashed lines, colored by subclone assignment; circles and squares denote non-synonymous and synonymous mutations, respectively. For driver mutations, the mutated gene and codon are indicated above each line. (B and D) Variability in intra-population growth rate, estimated by random sampling of 96 individuals at initial (*t* = 0 days, green) and final time points (*t* = 32 days,purple), before and after selection with (B) hydroxyurea and (D) rapamycin. Relative growth rates *λ*_k_(*t*) by individual *k* are shown at the foot of the histogram, calculated by averaging over *n*_*r*_ = 32 technical replicates per individual. Relative growth rates are normalized with respect to the mean population growth rate 〈*λ*_k_〉_*t*=0_ at *t* = 0 days. The posterior means of the distribution modes fitted by a Gaussian mixture model are indicated as dashed lines. The fitter individuals (pins) carry driver mutations, detected by targeted sampling and sequencing.The insets on the right-hand side depict a schematic of the fitness distribution in two limit cases: if there are many mutations of similar effect, the fitness wave will be smooth and unimodal; if only few mutations of large effect exist, the fitness distribution will become multimodal. See also Figures S3, S4 and S10.

### Selective Effects on Pre-existing Genetic Variation

To determine the adaptive value of background variation, we identified regions where local allele frequencies changed over the time course of the selection experiments. Frequency changes over time indicate that selection is acting on beneficial background alleles. These drivers cause linked passenger mutations to also change in frequency by genetic hitchhiking (Illingworth et al., 2012). We performed a systematic scan for background variants under selection using data up to 4 days, when no population yet had detectable subclones that would distort this signal (Supplemental Experimental Procedures). A region of interest was found in chromosome VIII (coordinates 460-490 kb) in all WAxNA populations under rapamycin (Figure 2B). We evaluated two candidate genes in this region by reciprocal hemizygosity, validating the *CTF8*^NA^ allele to increase rapamycin resistance. *CTF8* harbors two background missense variants and has previously been implicated in sensitivity to rapamycin, although the mechanism remains unknown (Parsons et al., 2004). Carrying the *CTF8*^NA^ allele confers a 36% growth rate advantage over the *CTF8*^WA^ allele (Figure S8). *KOG1*, which falls within the same region and is a subunit of the TORC1 complex, differs by seven missense mutations between the parents. However, reciprocal hemizygous deletions only revealed a modest fitness difference between WA and NA sequences of *KOG1*. We did not find events that replicated across all populations in hydroxyurea.

### Pervasive Selection of Macroscopic Subclones Driven by *De Novo* Genetic Variation

To reconstruct clonal expansions in the WAxNA populations we used background genetic variants as markers. Using the cloneHD algorithm (Fischer et al., 2014), we inferred the subclonal genotypes and their frequency in the populations, both of which are unknown *a priori* (Figure S1; Supplemental Experimental Procedures). We found at least one subclone in all WAxNA populations under selection, but none in the control environment (Figure 3 and S3). Clonal competition was prevalent with two or more expanding subclones in 12 out of 16 WAxNA populations. No population became fully clonal during the experiment, with subclone frequencies stabilizing after 16 days in several rapamycin populations. Similarly, WA and NA populations under selection underwent adaptation as evidenced by *de novo* mutation frequencies, except for WA which became extinct in hydroxyurea (Figure S4).

To genetically characterize the subclones, we isolated and sequenced 44 clones drawn from WAxNA populations after the selection phase (Figure 4; Experimental Procedures). From population and isolate sequence data, we observed 19 recurrent *de novo* mutations in the ribonucleotide reductase subunits *RNR2* and *RNR4* during hydroxyurea selection and in the rapamycin targets *FPR1* and *TOR1* during rapamycin selection (Table 1). Each of these driver mutations had a drug-resistant growth rate phenotype (Figures S6, S7 and S8) and carried a unique background of ~31,000 passenger mutations on average, compared to other sequenced isolates. All *FPR1* mutations were homozygous and likely to inactivate the gene or inhibit its expression. In contrast, *TOR1* mutations were heterozygous while we found *RNR2* and *RNR4* mutations in both heterozygous and homozygous state. All driver mutations occurred in highly conserved functional domains. The variant allele fractions of these mutations mirrored the inferred subclonal dynamics (Figure 3A, Figure 3C, S3 and S4). Other mutated genes with similar dynamics were confirmed as passengers (e.g. *DEP1*, *INP54* and *YNR066C*, see Figure S8). From the genome sequence of the 44 individual clones, we also detected six trisomies as large scale copy-number aberrations, without conclusive evidence that they are adaptive compared to recurrent point mutations (Figure 4).

**Figure 4:**
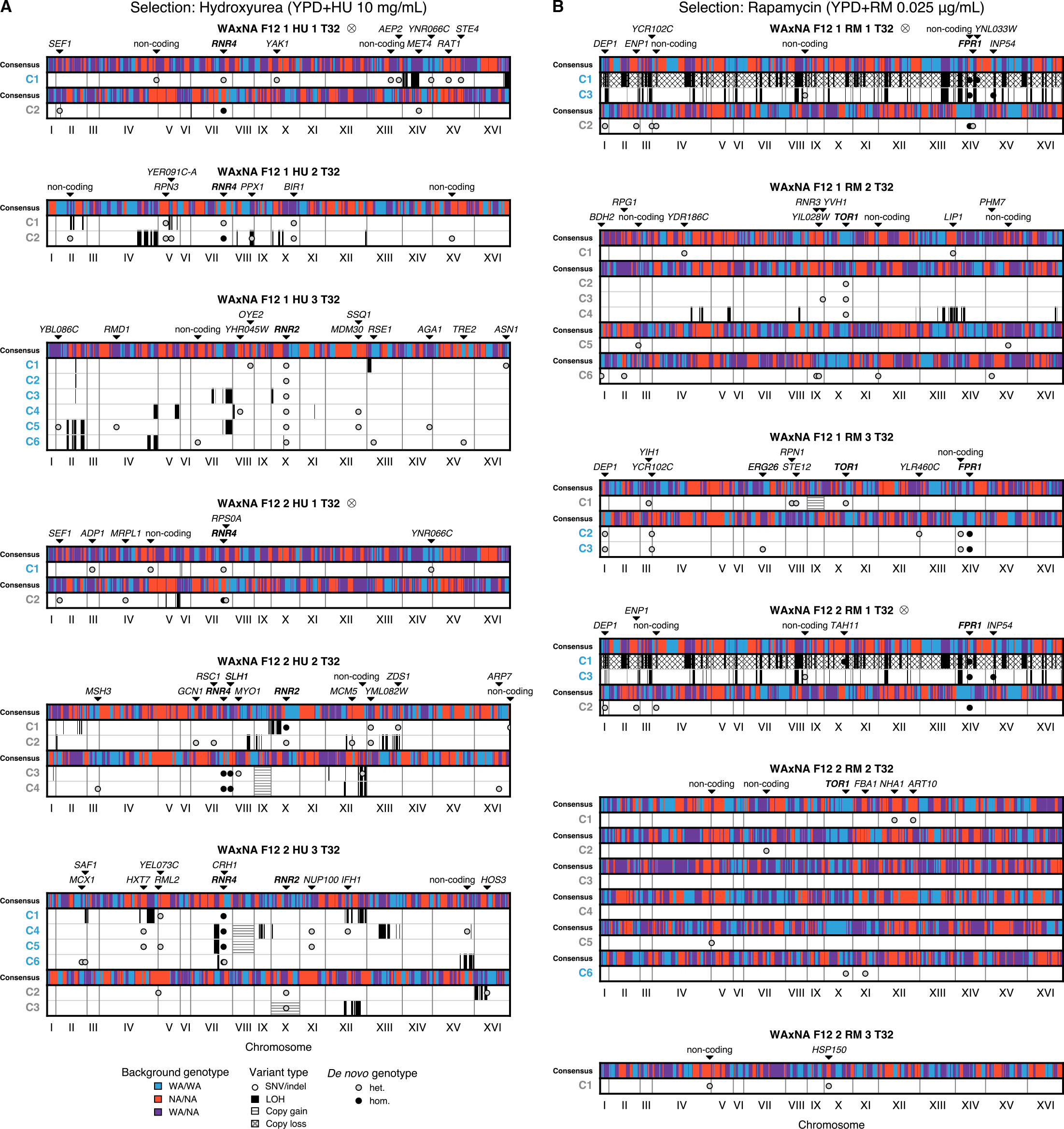
Pervasive Selection for Adaptive Mutations and Genomic Instability. Whole-genome sequences of clones sampled from WAxNA F_12_ populations. SNVs, indels and chromosome-level aberrations were detected by whole-genome sequencing in single-cell diploid clones derived from evolved populations, after *t* = 32 days in (A) hydroxyurea or (B) rapamycin(Table S1). Chromosomes are shown on the x-axis; clone isolates are listed on the left, colored by lineage (see Figure S3). The consensus shows the majority genotype across population isolates with sequence identity greater than 80%. WA/WA (in blue) and NA/NA (in red) represent homozygous diploid genotypes and WA/NA (in purple) represents a heterozygous genotype. Individual cells with shared background genotype carry *de novo* SNVs and indels (circles), *de novo* mis-segregations with loss-of-heterozygosity (solid segments) and *de novo* gains or losses in copy number (hatched segments). Driver and passenger mutations are listed along the top (drivers are in boldface). Populations marked by ⨂ indicate cross-contamination during the selection phase, but any derived events are independent. A figure with all ancestral sequenced isolates can be found in Figure S5. See also Figure 3A and Figure 3C, and Tables 1 and S1.

**Table 1.**
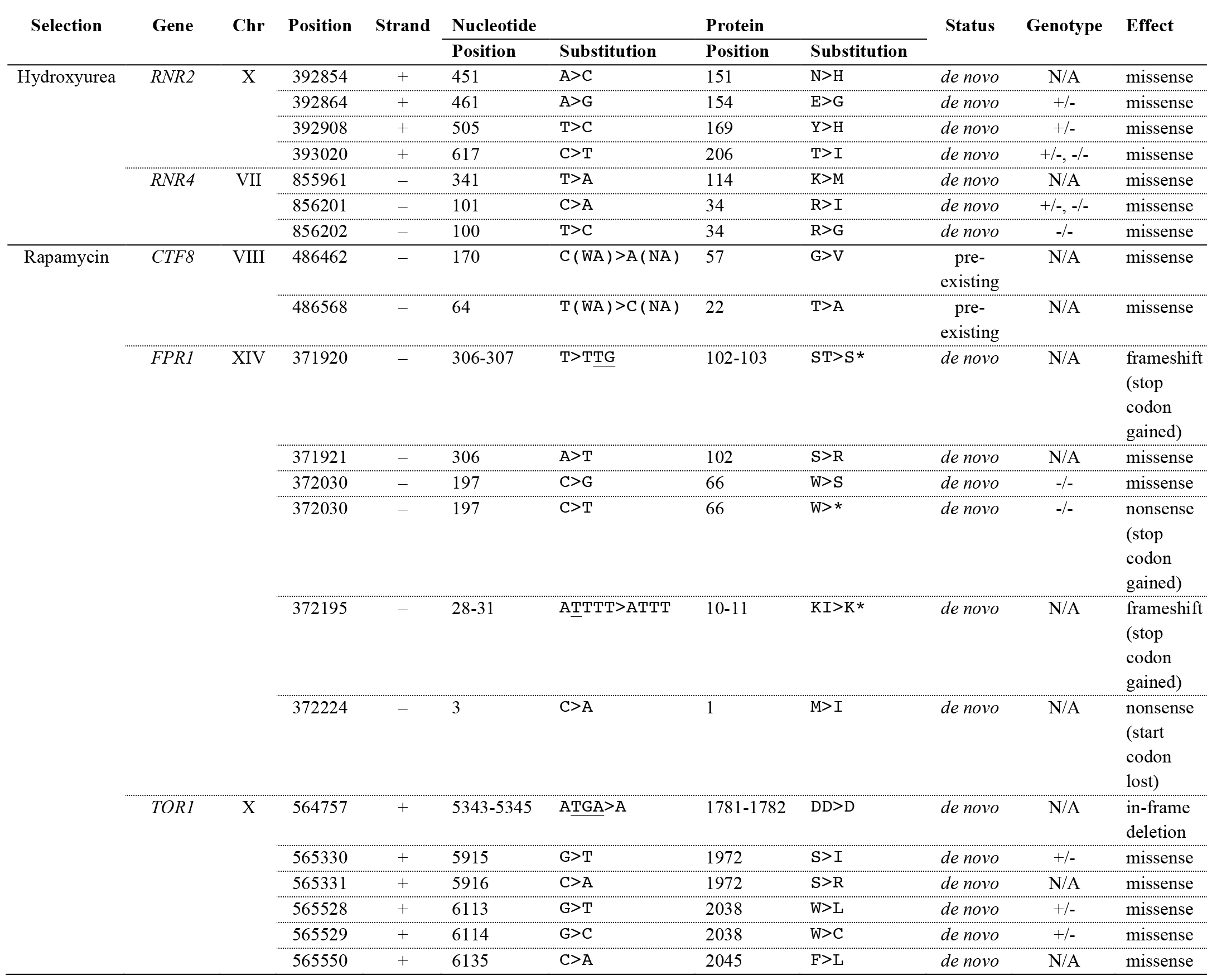
Summary of Driver Mutations. Summary of unique SNVs, insertions and deletions found to be drivers in hydroxyurea (*RNR2*, *RNR4*) and rapamycin (*CTF8*, *FPR1*, *TOR1*). Nucleotide and protein substitutions show the wild-type and mutated alleles. Nucleotides gained or lost are underlined. Variants are labeled as pre-existing if they differ between the parents, and *de novo* if they arose during the crossing or selection phases of the experiment. The functional impact of the mutations has been characterized using the Ensembl Variant Effect Predictor (McLaren et al., 2016). Populations and clones carrying mutations in these driver genes are listed in Table S1. The genotype of each mutation in individual clones is shown in Figure 4. The genotype of mutations only found by whole-population sequencing cannot be resolved, and is indicated as not applicable (N/A).

Clonal expansions were also evident from changes in the fitness distribution of cells. We established this by phenotyping 96 randomly isolated individuals from 3 populations per environment at 0 and 32 days, as well as the 44 sequenced individuals at 32 days (Experimental Procedures). We measured the growth rate of each isolate, and determined the population growth rate with respect to the mean of the fitness distribution. The variance of the fitness distribution varied significantly with different drugs, consistent with previous studies (Chevereau et al., 2015). While the variance of the fitness distribution at 0 days was narrow in hydroxyurea (σ^2^ = 3.1×10^−3^), growth in rapamycin showed a wider response (σ^2^ = 5.4×10^−3^). In rapamycin selection, the fitness distribution became multimodal after 32 days, reflecting the fitness of subclones substantially improving with respect to the mean fitness of the bulk population (Figure 3D). The clonal subpopulation divided on average twice as fast as the ancestral population. Sequenced isolates with driver mutations in *FPR1* and *TOR1* were on the leading edge of the fitness distribution, far ahead of the bulk. Furthermore, the bulk component showed a 10% average improvement, possibly due to selection of beneficial genetic backgrounds. Conversely, bimodality was only detected in one population in hydroxyurea selection (WAxNA F12 1 HU 3), where the clonal peak grew 25% faster on average compared to the ancestral, and the bulk 7% faster on average across all populations (Figure 3B). Isolates with *RNR2* driver mutations fell onto the leading edge of the fitness distribution. These six isolates originated from the same expanding subclone and two of them had 13% faster growth rate than the remaining four, although they all shared the same heterozygous *RNR2* driver mutation. In both of these clones, we found a large region in chromosome II to have undergone loss of heterozygosity (LOH), offering a putative genetic cause for their growth advantage (Figure 4A). Finally, to understand how the fitness of a typical population changes across environments we characterized the fitness correlations of ancestral and evolved clones, with and without stress (Figure S9). The rank order in clone fitness did not change significantly due to selection when measured in the absence of stress, implying that the evolutionary history of each of the clones did not lead to trade-offs in the average fitness of the population. However, a strong fitness cost of driver mutations in *FPR1* was observed.

### Diversification and Genomic Instability

We found several of the driver mutations to exist in homozygous rather than heterozygous states. LOH has been shown to rapidly convert beneficial heterozygous mutations to homozygosity in diploid yeast evolving under nystatin stress (Gerstein et al., 2014). Thus we hypothesized that genomic instability, causing widespread LOH, could be significantly contributing to adaptation. To detect mechanisms of genomic instability, we used heterozygous genetic variants as markers. Firstly, we used the sequences of haploid individuals from the ancestral population, drawn before the last round of crossing, to create *in silico* diploid genomes and calculate the length distribution of homozygous segments. Similarly, we measured the length distribution of homozygous segments from evolved isolate genomes. We observed a significant increase of long homozygosity tracts in the evolved clones - a hallmark of LOH (Figure 5A). Secondly, we directly counted LOH events in populations using multiple sequenced isolates from the same expanding subclone (Supplemental Experimental Procedures).

**Figure 5:**
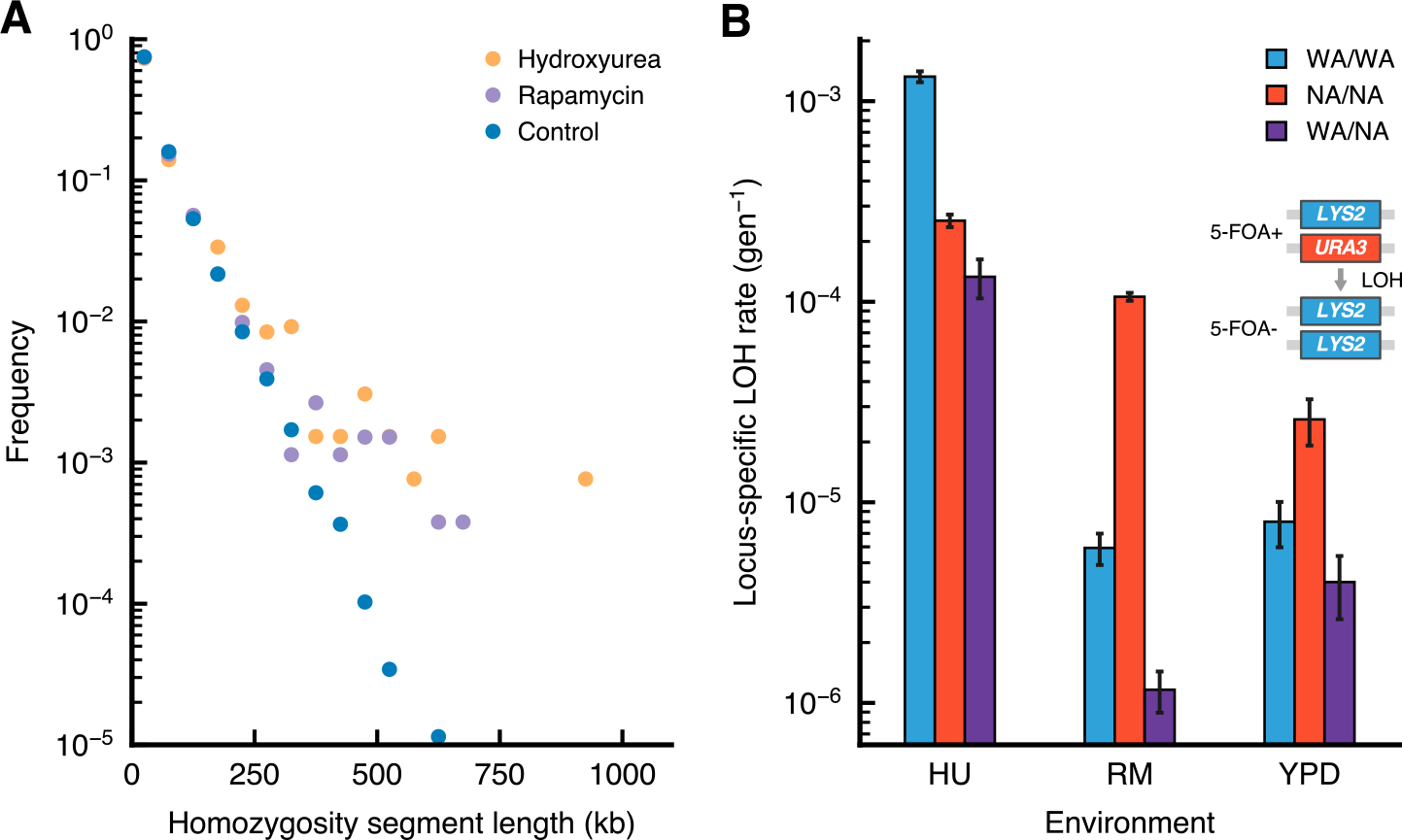
Elevated Rates of Loss of Heterozygosity. (A) The length distribution of homozygous segments, in bins corresponding to 50-kb increments, shows an excess of long homozygosity tracts above 300 kb in hydroxyurea and rapamycin (Kolmogorov-Smirnov test,*p* < 0.01). Ancestral haploid isolates are used to compare a set of *in silico* diploid genomes to evolved diploid isolates. Only unrelated isolate backgrounds were included. (B) Background-and environment-dependent rates of loss-of-heterozygosity were measured in a fluctuation assay by loss of the *URA3* marker. 5-FOA+ colonies indicate loss of the marker. Based on the number 5-FOA+ colony-forming units (CFU), the mean number of LOH events are estimated using the empirical probability generating function of the Luria-Delbrück distribution (Supplemental Experimental Procedures). The locus-specific LOH rates are shown, given by the mean number of LOH events divided by the total number of cells in YPD. Error bars denote the upper and lower 95% confidence intervals. LOH rates were elevated in hydroxyurea compared to the control environment and manifested background-dependent effects between the parents and their hybrid. See also Figure 4.

We identified a minimum of 6 events per genome per clone (Figure 4). Whilst this estimate is a lower bound and is limited due to the number of sequenced individuals per subclone, the LOH rates are substantial. To exemplify the interaction of genomic instability with pre-existing and *de novo* variation, inspection of *de novo* mutations in the WAxNA F12 1 HU 3 population shows that one *RNR2* mutation spans six isolates, being part of an expanding subclone (Figure 4A). These isolates have further diversified by acquiring passenger mutations and undergoing LOH. Clones C5 and C6 grow faster than the other four and share a large LOH event in chromosome II that is not present in the other isolates, possibly providing the growth advantage and broadening the fitness distribution (Figure 3B). An alternative route to homozygosity was observed in a single clone found to be haploid (clone C1 in WAxNA F12 2 RM 1) and therefore homozygous genome-wide. This haploid clone is closely related to a diploid clone (C3) from the same population and both clones share the same *FPR1 W66* de novo* mutation (Figure 4B). These data are consistent with the appearance of the FPR1 heterozygous mutation in an ancestral diploid clone that took two independent routes - focal LOH or meiosis - to unveil the recessive driver mutation. Altogether, we find that genomic instability can render *de novo* mutations homozygous as a necessary event in a multi-hit process towards drug resistance.

The stress environments themselves have an active role in accelerating genome evolution by chromosomal instability. Using a fluctuation assay, we investigated the effect of the genetic background and of the selective environment on chromosomal instability by tracking the loss of the *URA3* marker. Consistent with previous studies (Barbera and Petes, 2006), replication stress induced by hydroxyurea caused an increase in LOH rates. We also observed a background-dependent increase in LOH in rapamycin (Figure 5B).

### Decomposing Fitness Effects of Genetic Variation by Background Averaging

Finally, we sought to partition and quantify the individual fitness contributions of pre-existing and *de novo* genetic variation. The genotype space is extremely vast, but we can uniformly sample a representative ensemble to reconstruct a fraction of the genetic backgrounds where beneficial mutations could have arisen. To this end, we designed a genetic cross where background and *de novo* variants were re-shuffled to create new combinations (Figure 6A). We randomly isolated diploids from both ancestral and evolved populations, sporulated these and determined whether the derived haploids contained wild-type or mutated *RNR2*, *RNR4*, *FPR1* and *TOR1* alleles. We then crossed haploids to create a large array of diploid hybrids where all genotypes (+/+, +/-, -/-) for each of these genes existed in an ensemble of backgrounds, thus recreating a large fraction of the genotype space conditioned on the presence or absence of driver mutations. We measured the growth rates of both haploid spores and diploid hybrids, estimating and partitioning the variation in growth rate contributed by the background genotype and by *de novo* genotypes using a linear mixed model (Figure 6B and S10; Supplemental Experimental Procedures).

**Figure 6:**
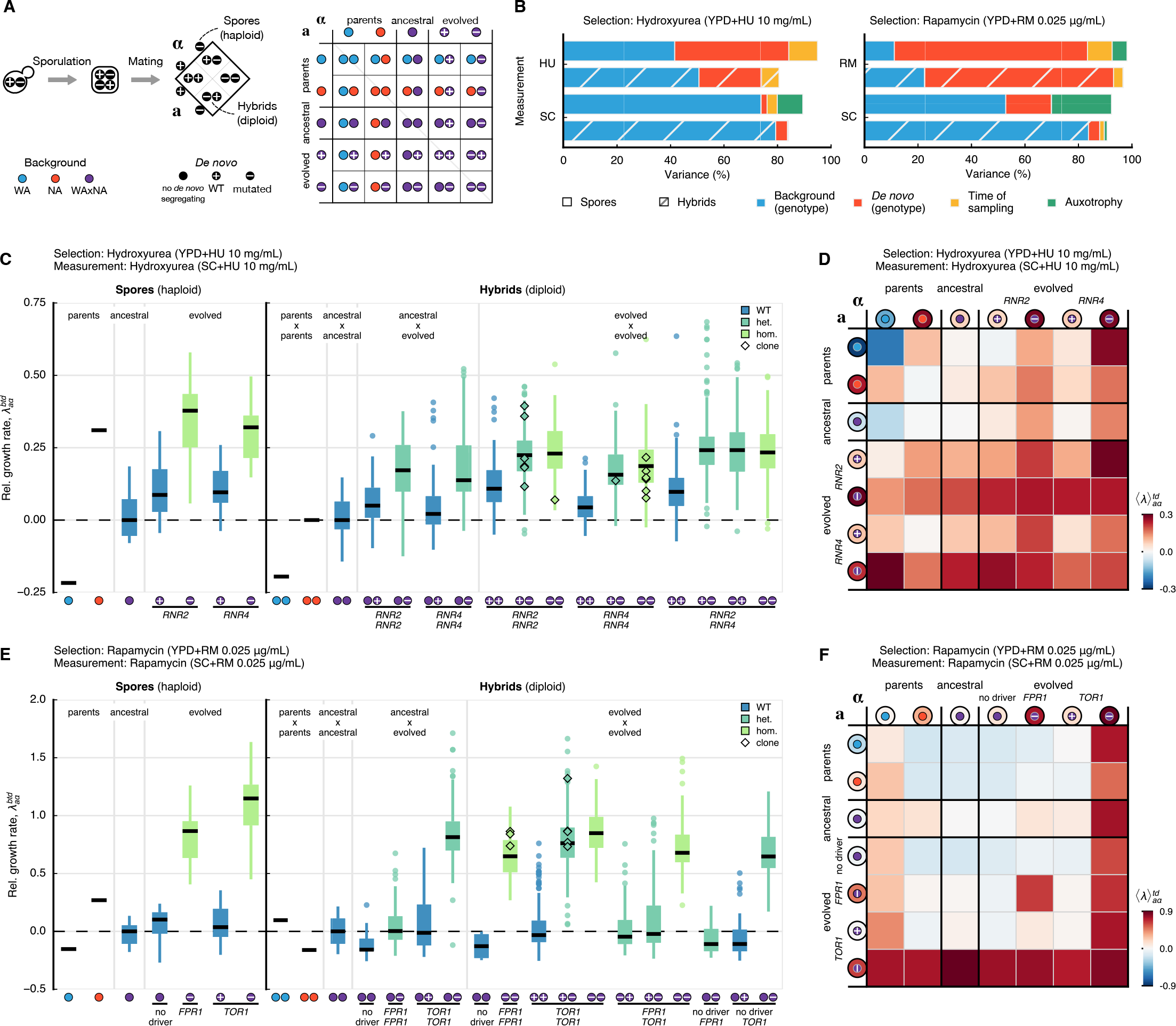
Ensemble-Averaged Fitness Effects of Genetic Background and *De Novo* Mutations. (A) To quantify the fitness effects of background variation and *de novo* mutations in hydroxyurea (*RNR2*, *RNR4*) and rapamycin (*FPR1*, *TOR1*), we isolated individuals from ancestral and evolved populations. From these diploid cells, we sporulated and selected haploid segregants of each mating type. Spores with mutations in *RNR2*, *RNR4* and *TOR1* were genotyped to test if they carry the wild-type or mutated allele. We crossed the *MAT*a and *MAT*a versions to create hybrids (48×48 in hydroxyurea and 56×56 in rapamycin). Independent segregants were used to measure biological variability of ancestral and evolved backgrounds. Symbols in panels C to F follow the legend in panel A and indicate combinations of the type of genetic background (WA parent, 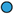; NA parent, 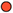; WAxNA segregant, 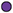) and the genotype of *de novo* mutations (no *de novo* mutation, 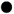; wild-type, 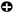 mutated, 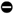). (B) Variance decomposition of the growth rate of spores (solid) and hybrids (hatched) that can be attributed to different components using a linear mixed model. The model components are the background genotype, b; *de novo* genotype, d; time of sampling during the selection phase, t; auxotrophy, x. Estimates of variance components are obtained by restricted maximum likelihood (Figure S12 and Table S6). (C and E) Relative growthrate of spores, 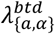, and hybrids, 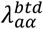, measured for multiple combinations of background and *de novo* genotypes and averaged over measurement replicates. Relative growth rates are normalized with respect to the mean growth rate of the ancestral WAxNA cross. Measurements of cells selected in (C) hydroxyurea and (E) rapamycin were taken in the respective stress environments. Medians and 25%/75% percentiles across groups are shown, with medians as horizontal black lines and colored by *de novo* genotype (wild-type, blue; heterozygote, cyan; homozygote, green). Outliers (circles) and isolated,selected clones with matching genotypes (diamonds) are highlighted. (D and F) Ensemble average of the growth rate of spores, 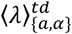, and hybrids, 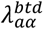 The color scale for all matrices is shown to the right of each panel and indicates the relative fold-change with respect to the ancestral WAxNA crosses.An extended version of the figure with all combinations and controls can be foundin Figures S10 and S11, respectively.

The ensemble average over backgrounds showed that the mean effect of *RNR2*, *RNR4* and *TOR1* mutations was fully dominant and highly penetrant regardless of the background (Figure 6D and Figure 6F). In contrast, *FPR1* mutants were recessive and only increased growth rate when homozygous, again irrespective of the background (Figure 6F). Recombinants with *RNR2* and *RNR4* mutations show epistatic interactions, consistent with the products encoded by these genes which are known to interact as subunits of the same protein complex (Figure 6C). After conditioning for *RNR2*, *RNR4*, *FPR1* and *TOR1* driver mutation status, a large fraction of the phenotypic variance still remained, reflecting the effect of the genetic backgrounds in which they emerged (Figure 6C and Figure 6E). In fact, under hydroxyurea exposure, background genetic variation accounted for an estimated 51% of the growth rate variance, more than twice the estimated 23% contributed by *RNR2* and *RNR4* de novo mutations. Furthermore, these mutations eventually landed on genetic backgrounds much fitter than average in the ancestral fitness distribution, as denoted by the estimated 7% explained by the time of sampling. Both of these results directly imply that moderate-effect de novo mutations must arise on favorable genetic backgrounds to give rise to macroscopic subclones. In contrast, under rapamycin exposure, the pre-existing genetic variation accounted for only 22% of the variance, much less than the 70% attributed to *FPR1* and *TOR1* mutations. Such large-effect mutations can expand in a vast majority of backgrounds, explaining how they can almost entirely surpass the bulk of the fitness distribution (Figure 3D). Taken together, these results are consistent with the aggregation of small-effect, pre-existing variants which can condition the fate of new mutations in both selection environments.

## Discussion

Here we showed that populations containing extensive fitness variability can adapt to strong selective pressures utilizing both pre-existing and *de novo* genetic variation. Theory predicts that pre-existing genetic variation forms a traveling fitness wave, with the mean fitness increasing at a rate that is proportional to its fitness variance (Desai and Fisher, 2007; Rouzine and Coffin, 2005). New mutations are expected to be successful if they land on a favorable background or if they are beneficial enough to escape from the bulk dynamics by their own merits. Recent theoretical results have suggested the existence of a selective advantage threshold above which the fate of a new mutation becomes decoupled from the background it lands on (Good et al., 2012; Schiffels et al., 2011). Our results show that new beneficial mutations expanded on a range of genetic backgrounds and selection concomitantly acted on pre-existing variation through its combined effects on fitness, steadily improving the bulk of the population. The observed dynamics are at this level consistent with the theoretical picture.

The rate of adaptation and what type of beneficial mutations are successful depend on multiple factors such as population size, mutation rate and ploidy (Barrick and Lenski, 2013; Selmecki et al., 2015; zur Wiesch et al., 2011). Our results show that sufficiently large populations could readily find beneficial *de novo* mutations, but their adaptive trajectories were simultaneously shaped by pree-xisting and *de novo* variation with overlapping timescales. Previous experimental studies with substantial founder diversity did not observe *de novo* mutations playing an important role in either asexual or sexual evolution (Burke et al., 2010, 2014; Parts et al., 2011). This may be due to differences in the selective constraints, which affect the timescale for the emergence of *de novo* mutations, or may depend on the genetic architecture of the selected phenotype making the background fitness variation harder to overcome. Despite the large genetic heterogeneity of the founders, mutations in driver genes were recurrent, indicating convergent evolution towards a restricted number of molecular targets. This is an important aspect to be able to predict the outcome of selection. Larger studies which systematically vary key parameters, such as population size, are needed to quantify how pre-existing variation modifies the reproducibility of new mutations.

Measurements of the fitness distribution revealed markedly different variability within a population in response to different inhibitors. There were two different outcomes of selection: if many mutations had comparable fitness effects as in hydroxyurea, the fitness distribution remained smooth; on the contrary when few large-effect mutations were available, such as mutations in the TOR pathway in rapamycin, the fitness distribution became multimodal. We were not able to attribute increases in the bulk of the fitness distribution to particular alleles beyond the *CTF8* gene, probably due to the contribution of many small-effect loci. Previous studies in isogenic populations have reported adaptive mutations sweeping to fixation in a comparable timescale, without specific selective constraints such as drugs (Lang et al., 2013). In contrast, we did not observe complete fixations. This is partially due to the duration of the experiment: the clones are still expanding after 32 days in hydroxyurea. However, most rapamycin-resistant clones become stable between 16 and 32 days. While we do not know the underlying cause, the observation has important consequences. Notably, the substantial genotypic and phenotypic diversity that remained after selection could be a potent substrate to re-sensitize a population, and may compromise targeted therapies against resistant clones. Understanding the role of clonal competition in isogenic and heterogeneous populations requires further work, which could be approached experimentally using lineage tracing (Levy et al., 2015).

We observed a balance between the loss of diversity due to selection, and active diversification mechanisms that partially re-established and refined existing variants. The background not only contributed substantially to fitness, but was also continuously re-configured by genomic instability, diversifying the expanding clones. Chromosomal rearrangements represent a key mechanism in shaping genome diversity in asexual organisms (Dunham et al., 2002; Flot et al., 2014) and in somatic evolution of cancer (Stephens et al., 2011), where cells accumulate a genetic load during tumor development that LOH can phenotypically reveal. In asexual diploids, such as studied here, successful beneficial mutations are expected to be dominant in a phenomenon known as Haldane's sieve (Orr and Betancourt, 2001). However, LOH has been shown to overcome this constraint by rapidly converting initially heterozygous mutations to homozygosity (Gerstein et al., 2014). Therefore LOH may enable asexually evolving populations to approach the adaptive rates seen in sexual organisms with recombination. Here we also saw these dynamics at play, as recessive *FPR1* mutations needed a second hit by LOH. Additionally, the process gained a new dimension: although these rearrangements were mostly copy-number neutral, they lead to fitness increments by changing scores of background variation from heterozygous to homozygous state in a single step. As a result, certain passenger mutations hitchhiking with a beneficial driver may provide an additional fitness advantage distributed across one or multiple loci (Figure 4). The implications of the ongoing diversification by chromosomal rearrangements are worthwhile pursuing further, both theoretically and experimentally. Even if a driver mutation fully fixed, a substantial amount of genetic variation would remain. Multiple genetic backgrounds with the same driver mutation will diverge (Hermisson and Pennings, 2005) and it may drastically alter the theoretical expectation of a sharp transition between evolutionary regimes at the selective threshold (Good et al., 2012; Schiffels et al., 2011). Experimentally, recently developed genome-editing techniques may enable localizing and measuring the fitness effect of specific LOH regions (Sadhu et al., 2016).

We carried out background-averaged fitness measurements of a recombinant library of pre-existing and *de novo* mutations. We found that large-effect mutations, such as those in the TOR pathway, confer resistance to rapamycin regardless of the genetic background where they arise. These mutations were of sufficient magnitude to surpass the bulk of the fitness distribution and can be interpreted to be above the selective threshold. Conversely, the pre-existing fitness variance influenced the fate of *de novo* drivers like *RNR2* and *RNR4* mutations, which needed to land on a favorable background to be competitive. Thus far, most biological systems have been found at the edge of the two regimes: large-effect mutations being amplified on well-adapted background genotypes have been observed in laboratory populations (Lang et al., 2011) and in the wild, e.g. in the seasonal influenza virus (Illingworth and Mustonen, 2012; Luksza and Lässig, 2014), which suggest that these dynamics represent a general mode of adaptation. Interestingly, our combinatorial strategy of background averaging shows that both of the limit cases can be true. Thus, predictability of the outcomes of selection will hinge on characterizing the background fitness variance and finding a common framework to describe the selective potential of a population (Boyer et al., 2016). Detecting a known driver mutation without a measurement of the background fitness distribution will be insufficient to predict its ultimate fate. This is a necessary requisite to eventually rationalize the design of therapies in the treatment of bacterial and viral infections or cancer. It may also be possible to balance and control the fitness effects of pre-existent and *de novo* mutations, i.e. to change the selective threshold, for example by modulating the dose-dependent effects of inhibitors (Chevereau et al., 2015) or by inhibiting global regulators (Jarosz and Lindquist, 2010).

Taken together, our findings can help to understand the evolution of large asexual populations with extensive genetic variation. Bacterial infections and cancer, which easily reach sizes of billions of cells, host a comparable mutation load before any selective treatment is applied. For example, the number of pre-existing variants in our experiment is comparable to the typical number of somatic mutations accrued before treatment during carcinogenesis, which varies between 10^2^-10^5^ depending on the cancer type (Lawrence et al., 2013); and it is also comparable to the genetic diversity in bacterial communities (e.g. in cystic fibrosis patients (Lieberman et al., 2014)). In either of these cases, the number of possible mutations available to escape antimicrobial or chemotherapy drugs is limited, and it is comparable to the balance we observe between the number of drivers and passengers. Clearly, whether or not these results hold more generally needs to be studied across systems. Overall, we hope our results will encourage new theoretical and empirical investigations of the complex interplay of selection simultaneously acting on pre-existing and *de novo* genetic variation, and of the role of genomic instability continuously molding the genomes in a population.

## Experimental Procedures

A summary of the experimental protocols of this study is presented in the Experimental Procedures. A full exposé of the experimental methods is given in the Supplemental Experimental Procedures, where we describe protocols for clone isolation, engineering genetic constructs, genetic crossing, fluctuation assays and growth phenotyping. This is followed by apresentation of the theory and data analysis, where we define the model for localization of drivers amongst hitchhiking passengers and the probabilistic inference method for subclonal reconstruction. Furthermore, we also discuss the model for the estimation of variance components from background-averaged fitness measurements. Supplementary Figures and Supplementary Tables are also appended in the Supplemental Information.

### Study Design

In our study, we begin with two yeast strains which have diverged over millions of generations (*divergence* phase), that are randomly mated by meiotic recombination to generate a large pool of recombinant mosaic haplotypes (*crossing* phase), followed by applying a selective constraint of the population under stress (*selection* phase).

#### Divergence phase

Parental strains were derived from a West African strain (DBVPG6044; *MATα*, *ura3::KanMX*, *lys2::URA3*, *ho::HphMX*) isolated from palm wine and a North American strain (YPS128; *MATa*, *ura3::KanMX*, *ho::HphMX*) isolated from oak tree. Hereafter we refer to these strains as WA and NA, respectively. These strains were selected from two diverged *S. cerevisiae* lineages and feature 52,466 single-nucleotide differences uniformly distributed across the genome.

#### Crossing phase

The selection experiments were carried out using WA, NA, WAxNA F_2_ and WAxNA F_12_ founder populations derived from hybrids between WA and NA. The WAxNA F_2_ and F_12_ populations were respectively generated from the F_1_ and F_11_ hybrids between WA and NA. The WAxNA F_1_/F_11_ diploid populations were expanded in YPD and sporulated in solid KAc medium (2% potassium acetate, 2% agar) for 14 days at 23°C. Sporulation of diploids was confirmed by visual inspection of asci. Over 90% of sporulation efficiency was observed after 14 days. Any remaining unsporulated cells were selectively removed using the ether protocol (Parts et al., 2011). The haploid population was subjected to mass mating according to the protocol described by Parts et al. (2011). Briefly, the asci were resuspended in 900 µl of sterile water and digested with 100 µl of zymolase (10 mg mL^−1^) for 1 hour at 37°C. The cells were washed twice with 800 µl of sterile water, vortexed for 5 minutes to allow spore dispersion, plated in YPD and incubated for 2 days at 23°C. The YPD plates were replica-plated in minimal medium to select diploid cells (*MATa/MATα*, *LYS2/lys2::URA3*). The WAxNA F_2_/F_12_ generation was collected from the plates and used as a founder population for the selection experiments as well as stored at −80°C as a frozen stock.

#### Selection phase

In the selection phase, WA, NA, WAxNA F_2_ and WAxNA F_12_ founder populations (referred to as *ancestral*) were evolved asexually in two selective environments and one control environment. Each of the ancestral populations consisted of a total population size of 3.2×10^7^ cells, determined by plating and counting colony-forming units. We serially propagated multiple replicate populations over a period of 32 days, which we refer to as *evolve*d populations. Every 48 hours, 1:10 of the total cell population was transferred to fresh plates, avoiding severe bottlenecks to minimize the impact of genetic drift. We estimated that 1.74 generations per day took place in hydroxyurea and 1.63 generations per day in rapamycin, based on the mean growth rate of three representative populations in each environment and accounting for acceleration and deceleration of growth every 48-hour cycle (Supplemental Experimental Procedures). These empirical estimates amount to ~54 generations between 0 and 32 days, in agreement with a theoretical bound on the number of generations assuming exponential growth with a 1:10 dilution factor every 48 hours.

Where indicated, the selective media were supplemented with hydroxyurea (HU) at 10 mg mL^−1^ or rapamycin (RM) at 0.025 µg mL^−1^ and maintained at constant drug concentration until day 34. The drug concentrations were chosen based on the dose response of the WA and NA strains. We selected concentrations that maximized the differential growth between the two diploid parents in each environment. We observed a clear dose response in hydroxyurea, with at least 10-fold differential growth between the two diploid parent strains at 10 mg mL-1 (Figure S7). For rapamycin we used 0.025µg mL^−1^, which also results in a 10-fold difference between the parent strains (Figure S8). This concentration is well below the minimum inhibitory concentration of 0.1 µg mL^−1^ originally used to identify the highly penetrant *TOR1* mutations in the lab strain (Heitman et al., 1991).

### Whole-Genome Sequencing and Phenotyping

We followed the evolution of these populations over the course of the experiment using whole-genome sequencing and phenotyping of the bulk population, of ancestral and of evolved isolates. WA and NA populations are labeled by their background, the environment in the selection phase and the selection replicate, e.g. NA RM 1. WAxNA populations are labeled by background, number of crossing rounds, cross replicate, selection environment and selection replicate, e.g. WAxNA F12 2 HU 1. Time series samples are labeled from T0 to T32 and isolate clones carry a suffix, e.g. C1, C2, etc. Whole-population sequencing was performed after *t* = 0, 2, 4, 8, 16 and 32 days, and ancestral and evolved individuals were also sequenced (Table S1). Genomic DNA was extracted from the samples using the Yeast MasterPure kit (Epicentre, USA). The samples were sequenced with Illumina TruSeq SBS v4 chemistry, using paired-end sequencing on Illumina HiSeq 2000/2500 at the Wellcome Trust Sanger Institute. Phenotyping of ancestral and evolved individuals was performed by monitoring growth after *t* = 0 and 32 days using transmissive scanning (Supplemental Experimental Procedures).

## Data and Software Availability

The study accessions for the sequence data reported in this paper are available from the European Nucleotide Archive (ENA) and the NCBI BioProject. The dataset in study accession PRJEB2608 corresponds to raw DNA sequence reads previously reported in Parts et al. (2011). The dataset in study accession PRJEB4645 corresponds to raw DNA sequence reads newly reported in this study. The dataset in study accession PRJEB13491 corresponds to mutation calls in the two aforementioned datasets. All datasets have been jointly analyzed in this manuscript.

Phenotype data, fluctuation assay data, code, and notebooks are available from the GitHub repository (https://github.com/ivazquez/clonal-heterogeneity).

## Author Contributions

1.V.-G.,J.W.,V.M. and G.L. designed research; F.S.,J.L.,B.B.,J.H.,A.B. and E.A.P. conducted experiments, I.V.-G.,A.F. and V.M. developed theory, implemented computational methods and analyzed data; and I.V.-G.,V.M. and G.L. wrote the paper.

## Acknowledgements

We thank Agnès Llored, Jordi Tronchoni and Martin Zackrisson for technical help; Elizabeth Gibson for support with library preparation and sequencing; and Erik Garrison, Daniel Kunz, Leopold Parts, David Posada and Magda Reis for critical reading of the manuscript. We also thank participants of the program on Evolution of Drug Resistance held at the Kavli Institute for Theoretical Physics (University of California, Santa Barbara) for discussions.I.V.-G. is a recipient of a Wellcome Trust PhD fellowship and a Sanger Early Career Innovation Award. This research was supported by the Wellcome Trust grants WT097678 (to I.V.-G.) and WT098051 (to V.M.), Fundación Ibercaja (to I. V.-G), ATIP-Avenir (CNRS/INSERM), Fondation ARC grant SFI20111203947, FP7-PEOPLE-2012-CIG grant 322035, French National Research Agency grant ANR-13-BSV6-0006-01, Cancéropôle PACA (AAP Emergence), and a DuPont Young Professor Award (to G.L.). F.S. was supported by ATIP-Avenir (CNRS/INSERM), Becas Chile grant 74130015, CONICYT/FONDECYT grant 3150156 and MN-FISB grant NC120043 postdoctoral fellowships. A.F. was supported by the German Research Foundation grant FI1882/1-1,J.L. by Fondation ARC grant PDF20140601375, B.B. by La Ligue Contre le Cancer grant GB-MA-CD-11287 andJ.H. by the French National Research Agency grant 11-LABX-0028-01.

